# Pathogenic *Leptospira* in dogs and rodents in Tha Wang Pha, Thailand – Prevalence, diversity and linked environments

**DOI:** 10.64898/2026.03.16.712015

**Authors:** Warirat Jaiwung, Théo Dokhelar, Serge Morand, Kittipong Chaisiri, Michel De Garine-Wichatitsky, Anamika Kritiyakan, Vanina Guernier-Cambert

## Abstract

Human leptospirosis is a disease of public health importance in Thailand, but the animal species involved in the transmission cycle have not been fully uncovered. This study investigated *Leptospira* infection in dogs and terrestrial micromammals in rural Nan Province, Thailand, and the pathogen diversity. Sera from 95 seemingly healthy dogs and kidney samples from 399 micromammals were analyzed using real-time PCR for *Leptospira* detection, followed by conventional PCR and sequencing of infecting *Leptospira*. We investigated environmental factors associated with *Leptospira* infection in micromammals, using data collected during trapping.

Real-time PCR revealed ongoing infection in 8.4% (8/95) of dogs and 10.0% (40/399) of terrestrial micromammals, with 12 infected species including *Bandicota indica, Berylmys berdmorei, Berylmys bowersi, Mus cervicolor, Mus cookii*, and *Hylomys suillus*. In this qPCR-positive micromammals, three pathogenic *Leptospira* species were identified: *L. interrogans, L. weilii*, and *L. borgpetersenii*. This represents the first confirmed detection of *L. weilii* in rodents in Thailand. Infected micromammals were found in agricultural and forest habitats but not in human settlements.

Our study demonstrates potential complex leptospirosis epidemiology in rural Thailand, with multiple species serving as pathogenic *Leptospira* reservoirs across diverse habitats, and some shared pathogen diversity with human leptospirosis cases in Thailand. Free-roaming dogs may serve as bridge hosts, transmitting zoonotic *Leptospira* from micromammals to humans by visiting both animal habitats and human settlements. These findings emphasize the need for integrated One Health surveillance approaches to control leptospirosis in rural communities.

## Introduction

Leptospirosis is a neglected tropical zoonotic infection. In Thailand, human leptospirosis remains a significant public health issue. Although the national incidence of human leptospirosis has dropped since a major outbreak in 1999-2003, the Epidemiological Surveillance Report 2023 published by the Department of Disease Control, Ministry of Public Health, reported that the incidence began to rise again in 2022, and in 2023 it reached a peak of 6.95 per 100,000 [1]. In terms of mortality and burden of disease, a study in 15 hospitals in the Sisaket province reported 32.7% of severe leptospirosis among 217 confirmed cases (2015-2018); 7% died at the time of admission, and 4 years after hospital discharge, 14% had died and 15% had developed severe kidney complication [2]. Infection is caused by direct or indirect contact with pathogenic *Leptospira*, which are maintained *via* chronic kidney infection in carrier animals. Chronic carriers as well as mammal reservoirs shed bacteria into the environment through their urine [3,4]. Bacterial infection is facilitated by skin abrasions or mucosal membrane contact. *Leptospira* infection is frequently correlated with environmental / climatic factors, e.g., flooding and heavy rainfall, particularly in north and northeastern Thailand [5], as well as the presence of animal reservoirs.

As common chronic carriers of pathogenic *Leptospira*, rodents have been investigated in Thailand. *Leptospira* seroprevalence in rodents trapped in four regions / 10 provinces in 1998 - 2000 was 4.8% (56/1164), with most seropositive rodents found northeast (7.1% seropositive overall; 12.5% in Buriram) [6]. A molecular study conducted in 2009 - 2010 reported *Leptospira* infection in 18%, 2% and 0% of rodents in Loei, Nan and Buriram provinces respectively, with *L. interrogans, L. borgpetersenii*, and *L. kirschneri* identified [7]. A study in northeastern Thailand (Buriram, Kalasin, Sisaket and Surin; 2011 - 2012) identified 3.6 % *Leptospira*-positive rodents, with *Bandicota indica, Rattus exulans*, and *R. rattus* being notable carriers [8]. Infecting *Leptospira* species were *L. interrogans, L. borgpetersenii*, and *L. wolffii*. Other animal species have been investigated as potential reservoirs of *Leptospira* strains infecting humans. A study in Nan province investigated *Leptospira* urinary shedding in cattle, pigs, dogs, and a goat, as well as asymptomatic humans [9]. All species were infected, prevalence varying between 8% and 12% in domestic mammals. A phylogenetic analysis revealed a cluster of *L. interrogans* (including 4 dogs, 3 pigs, 3 cattle, 1 human) and a cluster of *L. weilii* (including 13 cattle, 9 pigs, 2 dogs, 1 goat). *Leptospira weilii* had not been previously identified in Thailand. A few studies have targeted dogs specifically. In rural and urban areas in north, northeastern, and central Thailand, 4.4% of dogs (stray or client-owned) showed urinary shedding *versus* a 12% seroprevalence [10]. In Songkhla province, south Thailand, 32.4% (120/370) of stray dogs exhibited *Leptospira* antibodies against *L. interrogans* when using latex agglutination test, while 0.5% (2/370) tested PCR-positive [11]. Other studies were overwhelmingly serology surveys, showing various seroprevalence rates, e.g., 11% (23/210) dogs at Chiang Mai University small animal hospital [12] and 11% (6/55) dogs in Mahasarakham province [13].

Studies in Thailand have rarely explored the possible role of different animal carriers in human leptospirosis epidemiology, and the one study that explored human, cattle, pigs, dogs, a goat and the environment did not include rodents [9]. Furthermore, no study has explored the interaction between dogs and rodents and its potential impact on the transmission cycle. Studies have hypothesized that dogs could be a potential epidemiological bridge host involved in zoonotic transmissions between wildlife and humans [14]. The predominant free-roaming nature of rural dogs can increase their exposure to pathogens from a variety of sources, including *Leptospira* from infected rodents or wildlife prevalent in the natural and agricultural environments. Our study aimed at investigating *Leptospira* infection and diversity in dogs and terrestrial micromammals in rural villages in Nan province, Thailand. More specifically, we wanted to (i) assess pathogenic *Leptospira* infection and *Leptospira* diversity in domestic dogs and terrestrial micromammals and their possible role in relation to human leptospirosis, and (ii) identify high-risk landscapes in Nan province, Thailand.

## Material and Methods

### Ethics statement

Ethics permits were obtained from the Ethical Committees of Mahidol University or Kasetsart University, Bangkok, Thailand, for each relevant project from which we tested samples (CEROPath, BiodivHealthSEA, FutureHealthSEA, SEA-dog-SEA). All field activities were approved by the relevant authorities (such as the Department of National Parks or the Royal Forest Department Management) and with participation/supervision of local representatives. Informed consent was obtained from voluntary dog owners before sample and data collection.

### Sample collection of dogs and micromammals

Terrestrial micromammals were trapped between November 2012 and September 2018 around 8 villages in Tha Wang Pha district, Saen Thong Sub-district, Nan province: V1 Baan Nanoon, V2 Baan Nasai, V3 Baan Pho, V4 Baan Huak, V5 Baan Namkrai, V6 Baan Huaymuang, V7 Baan Santisuk and V8 Baan Hae (Figure 1) [15]. Dog samples were collected from 4 villages (V4, V5, V6, V7) in November 2019, October 2020, and January and July 2023.

**Figure 1.**
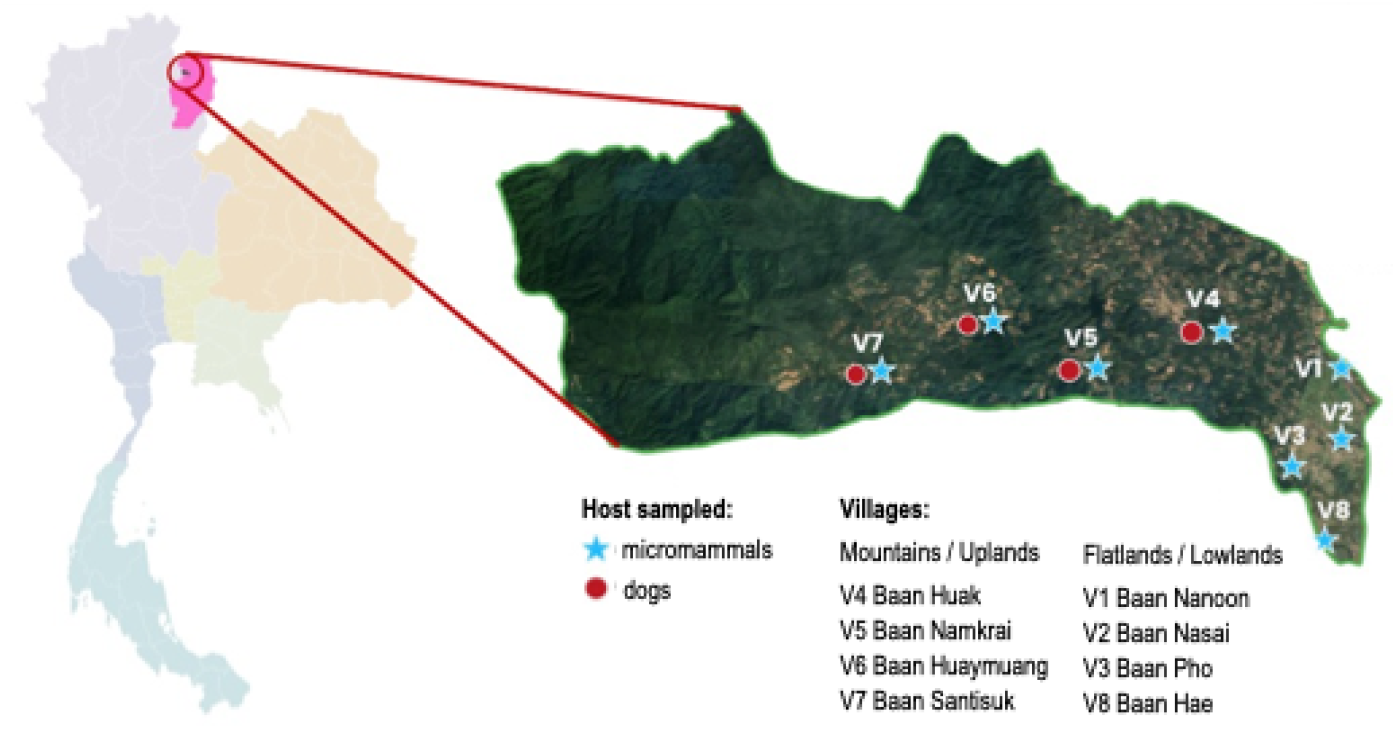
Study area in Nan province, Tha Wang Pha district, Saen Thong sub-district. Dogs were sampled in 4 villages; terrestrial micromammals were trapped in 8 villages.

Dogs were sampled as part of the FutureHealthSEA and the SEAdogSEA projects in November 2019, October 2020, January 2023, and July 2023. Healthy free-roaming adult dogs were selected after physical examination by a veterinarian and 5 mL blood was collected *via* venipuncture and stored at - 20°C. Terrestrial micromammals, defined as mammals of ≤1 kg live weight [16] were sampled as part of the CEROPath, BiodivHealthSEA, FutureHealthSEA projects, twice at each site: during the dry season (November 2012, March 2013, December 2013, February 2015 and November 2016) and the rainy season (July 2018, August 2018 and September 2018). Trapping involved the use of 30 lines of 10 traps (10 m apart) at each location for a four-night duration during each trapping session.

Information was collected for each individual, such as GPS coordinates, species, and landscape (divided in (1) forest and mature plantations, (2) upland (non-flooded lands), (3) lowland (cultivated floodplain), and (4) human settlement) [17]. Upland and lowland habitats were both agricultural lands (e.g., rice, corn) but were divided based on the geographic context, mountainous areas for “upland” vs flatlands for “lowland”. Sample collection followed the CERoPath protocol [18] and rodent species identification followed CERoPath application and barcoding tool (healthdeep.shinyapps.io/Small_mammals_CERoPath/). All micromammals were euthanized using an overdose of isoflurane in accordance with the guidelines of the American Society of Mammalogists and the European Union legislation (Directive 86/609/EEC). Subsequently, kidney samples were collected in 1.5 mL microtubes containing absolute ethanol and tubes were stored at -20°C.

### DNA extraction and pathogenic Leptospira detection

For dogs, DNA was extracted from 200 µl of dog serum using the NucleoSpin® Blood Mini Extraction Kit (Macherey-Nagel, Germany) following the manufacturer’s protocol, with a final elution volume of 80 µl per sample. For micromammals, kidneys were aseptically sliced to collect a cube of tissue approximately the size of a needle head (25 mg) at the interface of the cortex and medulla regions. This tissue was minced with a blade followed by lysis and DNA extraction using the DNeasy® Blood & Tissue Kits (QIAGEN Inc., Germany) or the QIAamp® DNA Mini Kit (QIAGEN Inc., Germany) following the manufacturer’s protocol, with a final elution volume of 100 µl per sample.

Previous studies have shown a better performance of reverse transcriptase real-time PCR (RT-qPCR) compared to qPCR for *Leptospira* detection [19–21]. *Leptospira* infection was assessed using a probe-specific RT-qPCR to detect a 87-bp region of rrs (16S) gene with the Lepto-F, Lepto-R primers and Lepto-S probe [22]. Amplifications were done using the iTaq® Universal Probes One-Step kit (Biorad, USA) using the following conditions: reverse transcription at 50°C for 10 minutes, initial denaturation at 95°C for 3 minutes, followed by 45 cycles of denaturation for 15 seconds at 95°C and annealing/extension for 15 seconds at 58°C. Samples with doubtful results (unusual PCR curve and/or very late Ct) were retested using the Quantinova® Probe PCR kit (QIAGEN, Germany) using the conditions: initial denaturation at 95°C for 2 minutes, and 45 cycles of denaturation for 5 seconds at 95°C and annealing/extension for 5 seconds at 60°C. A *Leptospira*-positive sample was defined as a sample that either had a clear positive One Step qPCR (not retested) or had two positive qPCR results, whatever the Ct values. All qPCRs were performed on a CFX Opus96 thermocycler (BioRad, Germany).

### Leptospira genotyping

*Leptospira*-positive samples were tentatively amplified by conventional PCR using 3 pairs of primers. First, 527-bp fragments were amplified using the primers secYFd / secYR3 [23]. When *secY* PCR amplification failed, sequencing of alternative regions was performed: a 541-bp sequence from the *rrs* (16S) gene using primers rrs2F / rrs2R [24], and if necessary a 335-bp sequence from the *lfb1* gene using primers lfb1-F / lfb1-R [25].

PCR amplifications were performed in 20 µl containing 2 µl of nucleic acids template, 10 µl of 2x HotStarTaq (Plus) Master mix, 1 µl (0.5 µM) of each primer, and 4 µl of RNase-free water, using either the QIAGEN**®** HotstarTaq Plus Master Mix (QIAGEN Inc., Germany; discontinued during our study) or the QIAGEN**®** HotstarTaq Master Mix (QIAGEN Inc., Germany) using the following conditions: 15 minutes of initial activation at 95°C and 45 cycles of denaturation for 30 seconds at 94°C, annealing for 30 seconds at 55°C (*secY* gene), 58°C (*rrs* gene) or 57°C (*lfb1* gene), extension for 1 minute at 72°C and 1 cycle of final extension for 10 minutes at 72°C using a T100 Thermal Cycler (Bio-Rad, Germany). PCR products were run on a 1.5% gel at 100 V for 30 minutes, and amplicons matching the expected size were sent to Macrogen (South Korea) for Sanger sequencing.

### Data analyses

Sanger sequences were cleaned, and consensus sequences and alignments were performed using Geneious Prime software version 2025.2 [26]. Phylogenetic trees were constructed based on the maximum-likelihood (ML) method with 1,000 bootstraps, using PhyML 3.0 [27]. Trees were visualized in FigTree v1.4.4 (http://tree.bio.ecd.ac.uk/). GenBank accession numbers of the sequences produced during our study are provided as supplementary information (S1 Table). Statistical analyses were conducted in R version 4.5.2 [28]. Descriptive statistics and stratified summaries were generated with the table1 package version 1.4.3 [29].

We assessed the effect of individual and environmental factors on *Leptospira* infection in micromammals using a generalized linear mixed model (GLMM) with a binomial error distribution and a logit link. Infection status was designated as the response variable, while landscape, season, sex, standardized body mass and host species were fixed effects. We included year and village * year interaction as random effects to account for the non-independence of data due to spatio-temporal clustering. Season was coded as a binary factor (wet *vs* dry) based on the month of trapping following standard climatological definitions in Thailand, i.e., wet season defined as May to October and dry season as November to April. To account for within-species variation in body mass, we standardized all body masses to a species-specific Z-score prior to modelling. To ensure sufficient sample size per taxon and avoid model instability, analyses were limited to species with at least five individuals. The model was fitted in R version 4.5.2 using the glmmTMB package version 1.1.14 [30] with a binomial family and logit link. Model-based predicted infection probabilities and 95% confidence intervals were obtained using the ggeffects package version 2.2.3 [31]. Predicted probabilities for the body mass effect were visualized as a function of standardized body mass.

## Results

### Animal sampling

In total we collected sera from 95 owned domestic dogs from 4 villages: 25 from V4 Baan Huak, 24 from V5 Baan Namkrai, 27 from V6 Baan Huaymuang, 19 from V7 Baan Santisuk. We also collected 399 terrestrial micromammals from 13 rodent species, and one gymnure species (*Hylomys suillus*) (not counting unidentified *Berylmys* and *Mus* sp.; see Table 1). The most commonly trapped species were *Rattus exulans* (28%) followed by *Bandicota indica* (12%), *Mus cervicolor* (10%) and *Maxomys surifer* (10%). Micromammals were trapped in four habitats: human settlements (33%), lowland (26%), forests (22%) and upland (19%). Despite equal sampling effort, sampling success strongly varied between habitats and villages. Eight of 14 species were not trapped in human settlements, with *Rattus exulans* representing the majority (83%) of individuals trapped in this habitat. All 14 species were identified in V6 (where 72% of micromammals were trapped), while richness was 3-4 species in the other villages (where 2% to 7.5% of micromammals were trapped).

**Table 1.** Sample size of trapped terrestrial micromammals and *Leptospira* qPCR-positive per species. For each host species, the first line indicates the ratio of individuals of this species to individuals of all species per landscape (percentage); the second line (in italic) indicates the ratio of *Leptospira* qPCR-positive individuals to the total number of individuals of this species trapped in this specific landscape (percentage). One *Rattus tanezumi* (out of 32) which landscape was missing was excluded; 398 total individuals was used for prevalence calculations instead of 399.

| Host species | Forest<br>x/88 (%)<br>*pos/x (%)* | Lowland<br>x/102 (%)<br>*pos/x (%)* | Settlement<br>x/133 (%)<br>*pos/x (%)* | Upland<br>x/75 (%)<br>*pos/x (%)* | All<br>x/398 (%)<br>*pos/x (%)* |
| --- | --- | --- | --- | --- | --- |
| *Bandicota indica* | 1/88 (1.1)<br>*0/1 (0)* | 34/102 (33.3)<br>*3/34 (8.8)* | 0/133 (0)<br>*-* | 12/75 (16.0)<br>*3/12 (25)* | 47/398 (11.8)<br>*8/47 (17)* |
| *Berylmys berdmorei* | 4/88 (4.5)<br>*1/4 (25)* | 1/102 (1.0)<br>*1/1 (100)* | 0/133 (0)<br>*-* | 6/75 (8.0)<br>*1/6 (16.7)* | 11/398 (2.8)<br>*3/11 (27.3)* |
| *Berylmys bowersi* | 5/88 (5.7)<br>*2/5 (40)* | 6/102 (5.9)<br>*1/6 (16.7)* | 0/133 (0)<br>*-* | 14/75 (18.7)<br>*2/14 (14.3)* | 25/398 (6.3)<br>*5/25 (20.0)* |
| *Berylmys* spp. | 0/88 (0)<br>*-* | 1/102 (1.0)<br>*0/1 (0)* | 0/133 (0)<br>*-* | 2/75 (2.7)<br>*0/2 (0)* | 3/398 (0.8)<br>*0/3 (0)* |
| *Hylomys suillus* | 1/88 (1.1)<br>*0/1 (0)* | 0/102 (0)<br>*-* | 0/133 (0)<br>*-* | 5/75 (6.7)<br>*1/5 (20)* | 6/398 (1.5)<br>*1/6 (16.7)* |
| *Leopoldamys herberti* | 3/88 (3.4)<br>*0/3 (0)* | 1/102 (1.0)<br>*0/1 (0)* | 0/133 (0)<br>*-* | 0/75 (0)<br>*-* | 4/398 (1)<br>*0/4 (0)* |
| *Maxomys surifer* | 27/88 (30.7)<br>*1/27 (3.7)* | 2/102 (2.0)<br>*1/2 (50)* | 0/133 (0)<br>*-* | 11/75 (14.7)<br>*0/11 (0)* | 40/398 (10.1)<br>*2/40 (5)* |
| *Mus caroli* | 0/88 (0)<br>*-* | 4/102 (3.9)<br>*0/4 (0)* | 0/133 (0)<br>*-* | 1/75 (1.3)<br>*0/1 (0)* | 5/398 (1.3)<br>*0/5 (0)* |
| *Mus cervicolor* | 3/88 (3.4)<br>*0/3 (0)* | 31/102 (30.4)<br>*4/31 (12.9)* | 2/133 (1.5)<br>*0/2 (0)* | 5/75 (6.7)<br>*2/5 (40)* | 41/398 (10.3)<br>*6/41 (14.6)* |
| *Mus cookii* | 0/88 (0)<br>*-* | 8/102 (7.8)<br>*2/8 (25)* | 7/133 (5.3)<br>*1/7 (14.3)* | 2/75 (2.7)<br>*0/2 (0)* | 17/398 (4.3)<br>*3/17 (17.6)* |
| *Mus pahari* | 4/88 (4.5)<br>1/4 (25) | 0/102 (0)<br>- | 0/133 (0)<br>- | 0/75 (0)<br>- | 5/398 (1.3)<br>1/4 (25) |
| *Mus spp.* | 0/88 (0)<br>- | 2/102 (1.9)<br>0/2 (0) | 0/133 (0)<br>- | 0/75 (0)<br>- | 2/398 (0.5)<br>0/2 (0) |
| *Niviventer mekongis* | 24/88 (27.3)<br>0/24 (0) | 2/102 (2.0)<br>0/2 (0) | 0/133 (0)<br>- | 7/75 (9.3)<br>1/7 (14.3) | 33/398 (8.3)<br>1/33 (3.0) |
| *Rattus andamanensis* | 13/88 (14.8)<br>1/13 (7.7) | 0/102 (0)<br>- | 1/133 (0.8)<br>0/1 (0) | 2/75 (2.7)<br>0/2 (0) | 16/398 (4)<br>1/16 (6.3) |
| *Rattus exulans* | 1/88 (1.1)<br>0/1 (0) | 1/102 (1.0)<br>0/1 (0) | 110/133 (82.7)<br>8/110 (7.3) | 1/75 (1.3)<br>0/1 (0) | 113/398 (28.4)<br>8/113 (7.1) |
| *Rattus tanezumi* | 2/88 (2.3)<br>0/2 (0) | 9/102 (3.9)<br>1/9 (11.1) | 13/133 (9.0)<br>0/13 (0) | 7/75 (5.3)<br>0/7 (0) | 31/398 (7.8)<br>0/31 (0) |
| All species | 88/398 (22.1) | 102/398 (25.6) | 133/398 (33.4) | 75/398 (18.8) | 398/398 (100) |

### Leptospira detection and prevalence in animal specimens

Overall, we identified *Leptospira* DNA in 40 out of 399 (10%) micromammals, and in 12 out of 15 micromammal species (Table 1). Among *Leptospira*-infected micromammals, *Bandicota indica* (8/40) and *Rattus exulans* (8/40) were the most common species, each accounting for 20% of all infected. However, when looking at the *Leptospira* prevalence of each species, *Berylmys berdmorei* showed the highest infection rate (27%), followed by *Berylmys bowersi* (20%), *Bandicota indica* (17%) and *Hylomys suillus* (17%) (Table 1). *Leptospira* qPCR-positive micromammals were collected in 6 out of 8 villages: V2 (2.5%), V3 (2.5%), V5 (5%), V6 (77.5%), V7 (7.5%) and V8 (5%) (S2 Table). In village V6, 11 out of 15 species showed some *Leptospira*-positive individuals. In dogs, 8 out of 95 sera (8.4%) tested qPCR-positive for pathogenic *Leptospira*. They originated from three out of four villages: 2/25 (8%) dogs in V4 Baan Huak, 2/24 dogs (8.3%) in V5 Baan Namkrai, 0/27 in V6 Baan Huaymuang and 4/19 dogs (21%) in V7 Baan Santisuk.

### Leptospira genetic diversity in animals from Nan and in Thailand human cases

We successfully identified the infecting *Leptospira* in 11 out of 40 (27.5%) qPCR-positive micromammals, with three pathogenic species identified: *L. interrogans* (n = 5), *L. borgpetersenii* (n = 2) and *L. weilii* (n = 4). We were not able to either amplify or sequence the infecting *Leptospira* for the eight infected dogs. We built *secY*-based (Figure 2) and *rrs*-based (S1 Figure) phylogenetic trees including 11 and 7 sequences from our micromammals respectively. We also included published sequences from GenBank from Thailand and other Southeast Asian countries, originating from different host species, including dogs. Our *secY*-based phylogenetic analysis revealed that R6916 (*B. bowersi*), R7230 (*B. indica*), R7157 (*H. suillus*), R7572 (*B. bowersi*) and R6775 (*B. berdmorei*) clustered within the *L. interrogans* clade. They were genetically closely related to dog, human and rodent *Leptospira* sequences from Thailand and Cambodia. R7698 and R7150 (from *B. indica*), R7189 (*B. bowersi*) and R7599 (*B. berdmorei*) clustered with *L. weilii* strains isolated from dogs and humans *Leptospira* sequences from Thailand. R7827 (*Mus cookii*) and R6928 (*Mus cervicolor*) clustered with *L. borgpetersenii* isolates from humans and rodents from Thailand and Cambodia. While *L. borgpetersenii* was not identified in dogs using *secY* primers, published *rrs* sequences showed that dogs can be infected with *L. borgpetersenii*.

**Figure 2.**
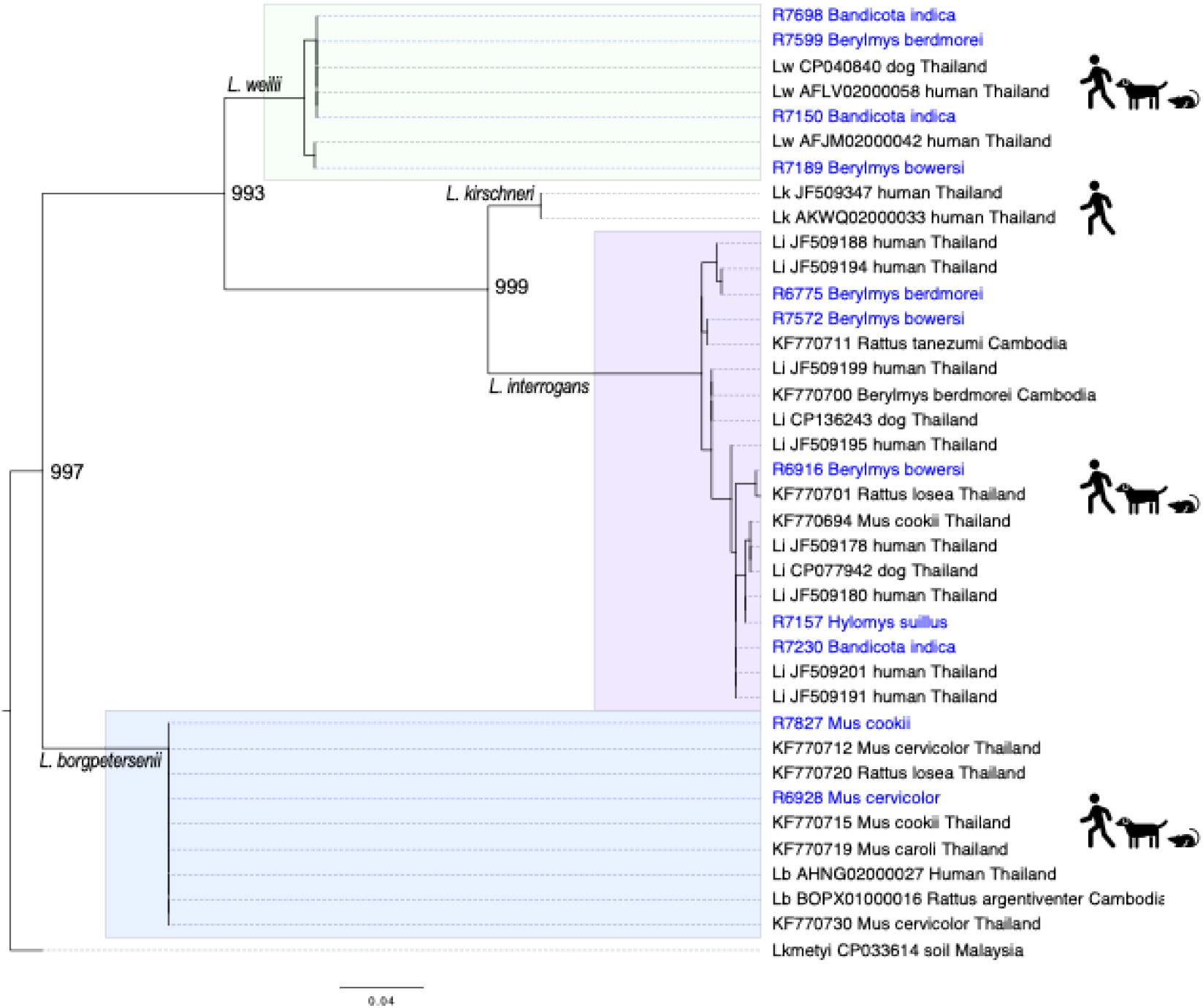
Maximum-likelihood phylogenetic tree (model HKY85; 1,000 replicates) inferred from *secY* gene (459-bp sequence). Samples from our study (n = 11) are in blue, using identifiers accompanied by the host species. Published sequences (in black) are identified using *Leptospira* species followed by the GenBank accession numbers, the source/host species and the country of origin. Colored silhouette icons indicate the host families identified within each clade associated with a specific *Leptospira* species. Bootstrap values are indicated for the main nodes.

### Explanatory model for micromammals infection status

The binomial GLMM revealed no statistically significant association between landscape type and *Leptospira* infection in terrestrial micromammals after accounting for random effects of trapping year and village * year interaction (S3 Table). Although estimated infection risk tended to be higher in lowland and upland habitats relative to forest, none of these differences were statistically supported (lowland: z = 1.33, p = 0.18; upland: z = 0.72, p = 0.47; settlement: z = 0.24, p = 0.81). Predicted infection probabilities were generally low across all landscape categories, with wide and largely overlapping 95% confidence intervals, providing no clear evidence that landscape type influenced *Leptospira* infection risk in micromammals.

Season had no detectable effect on infection probability (z = −0.56, p = 0.58), nor did sex (z = −0.82, p = 0.41). In contrast, standardized body mass showed a positive significant association with infection probability (z = 2.20, p = 0.03), indicating that individuals heavier than their species average tended to have a higher probability of infection (Figure 3).

**Figure 3.**
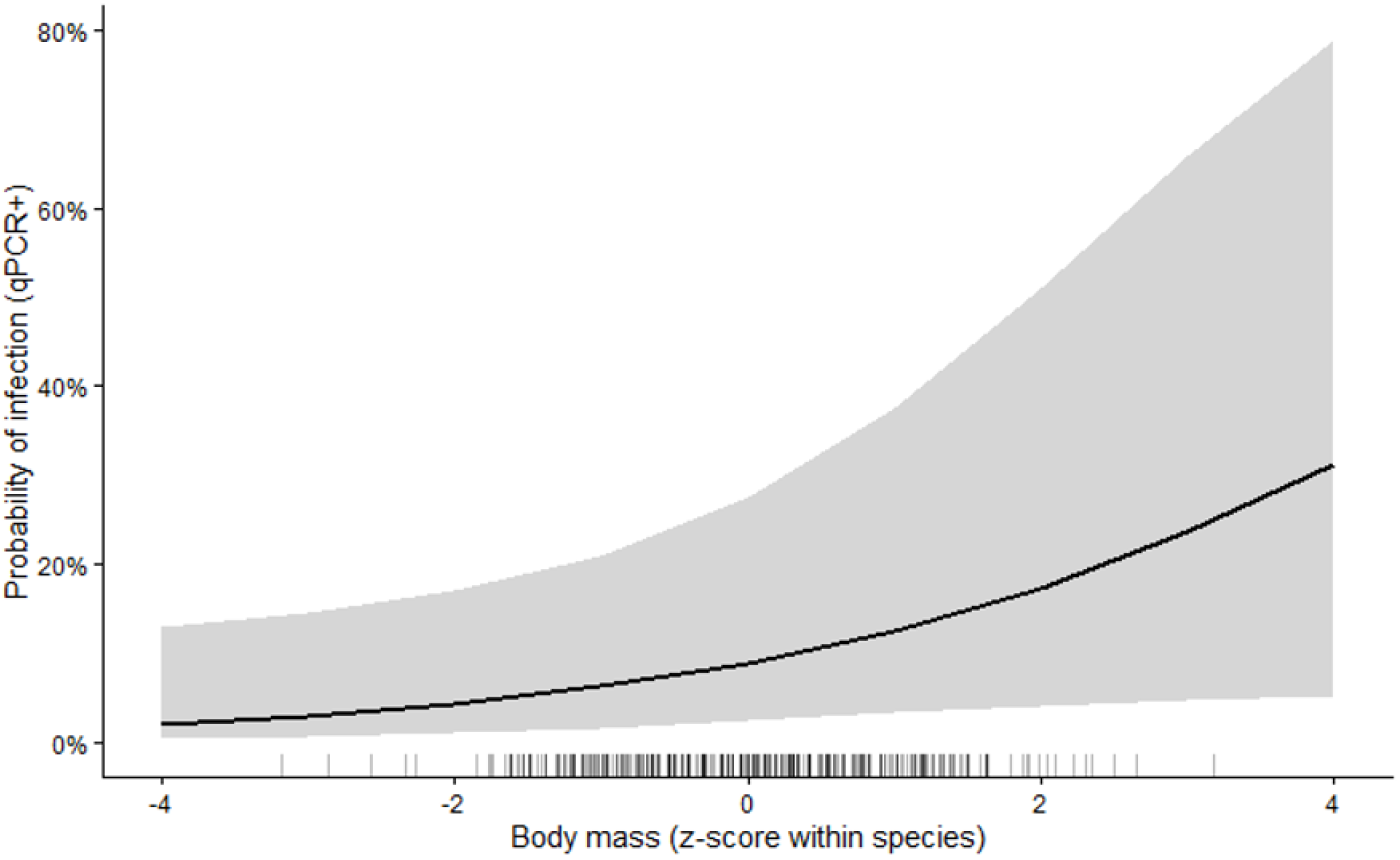
Probability of *Leptospira* infection in terrestrial micromammals as a function of standardized body mass. Body masses were standardized to a species-specific Z-score. The solid line represents binomial GLMM predictions of the probability of *Leptospira* infection, while the shaded areas indicate 95% confidence intervals. Rugs along the x-axis indicate the distribution of standardized body masses. Predictions were adjusted for landscape, season, sex, and host species with random intercepts accounting for year and village * year interaction.

Random effects indicated negligible variation attributable to year alone (SD ≈ 0) but substantial variation attributable to the village * year interaction (SD = 0.34 on the logit scale).

## Discussion

Our study in Saen Thong subdistrict, Nan province, confirms that multiple terrestrial micromammals (12 infected species in our study) serve as natural reservoirs for pathogenic *Leptospira* in the study area, with an overall 10% infection rate over the period 2012-2018. Among all the infected, *Bandicota indica* and *Rattus exulans* were the most common species. When considering each species separately, *Leptospira* prevalence was highest (17 to 27%) in *Berylmys berdmorei, Berylmys bowersi, Bandicota indica*, and *Hylomys suillus*. In a previous study conducted in Nan province (2008-2009), 4.4% (10/225) rodents were found infected with *Leptospira*, including *Bandicota indica, Berylmys berdmorei* and *Berylmys bowersi* while *Mus cervicolor* and *Mus cookii* consistently tested negative [32]. In another study, rodent samples collected from Loei, Nan and Buriram (2009-2010) showed various *Leptospira* prevalence (18%, 2% and 0% respectively) [7]. *Mus cookii* and *Berylmys berdmorei* had the highest prevalences, but other species (e.g., *Bandicota indica, Maxomys surifer* and several species of rats and mice) were also infected. Our study confirms other observations in Thailand that terrestrial micromammals carrying pathogenic *Leptospira* are highly diverse, even in a small area [7,33].

In upland Saen Thong, 8.4% (8/95) sera from apparently healthy dogs tested *Leptospira*-positive. In a multi-species study conducted in Nan province in 2013-2016, *Leptospira* urine shedding was 10.3% (6/58) in healthy dogs [9]. In a larger study that tested rural and urban dogs from northern, northeastern, and central Thailand, *Leptospira* urine shedding was 4.4% (12/273) [10]. In Southern Thailand, researchers found that 0.5% of healthy stray dogs tested in 2014-2018 were infected with *Leptospira* but 32.4% were seropositive, indicating a low infection rate at the time of collection but high levels of past exposure in the stray population [11]. In a study conducted in 2020-2023 through a neutering program in seven provinces across the central, western and southern regions, *Leptospira* urine shedding was 11.2% (34/303) [34]; the infection frequency in owned dogs (10.2%) was lower than that observed in not-owned (free-roaming) dogs (11.8%) but the difference was not statistically significant. While infection rates may vary between provinces and dog populations, the detection of pathogenic *Leptospira* DNA in the blood or urine of apparently healthy dogs is a public health concern as those dogs can shed pathogenic *Leptospira*, contaminating the environment [35]. This probably contributes to the chain of transmission between dogs but also possibly between species, including humans.

In our study, standardized body mass emerged as the main host-related predictor of *Leptospira* infection. Individuals heavier than their species average had a significant higher probability of infection, suggesting that the probability of infection may increase with age and/or body condition. Similar positive associations between rodents body mass and *Leptospira* infection have been reported in several studies, where larger older individuals exhibited higher infection prevalence [36,37]. Although ‘host species’ was included as a variable in the model, species-level differences in the probability of infection were associated with uncertainty, partly due to low numbers of infected individuals in some taxa. Environmental factors showed limited influence in our model. Infection probability exhibited minor variation across landscapes, with a non-significant tendency toward higher prevalence in lowland and upland compared to forest and human settlements. Season and sex also had no detectable effect on infection probability. Of note, despite equal sampling effort, 72% of terrestrial micromammals were trapped in village 6, which might have introduced a bias in our risk analysis regarding individual and environmental factors. Such spatial imbalance could have reduced our power to detect village- and landscape-related effects. In a previous study in Nan province, Thailand, elevated leptospirosis risk for humans (2003-2012) was documented in open habitat near rivers and in rice fields prone to flooding, while infected rodents (collected in 2008-2009) were more frequently found in patchy habitats with high forest cover, mostly situated on sloping ground areas [32]. In a later study conducted in several Thai provinces, the main risk factors for developing severe human leptospirosis were living near a rubber plantation and bathing in natural bodies of water [38]. These heterogenous findings indicate no clear pattern, suggesting that leptospirosis epidemiology in Thailand is complex, and that local epidemiology may vary between sites, with different animal reservoirs and associated risk factors possibly involved. In villages 4, 5, 6 and 7, a survey of dog owners identified that the majority of dog owners living in the upland forested areas tended to keep their dogs outside houses at all time [39], which can increase their likelihood of coming into contact with high-risk environments and diverse reservoirs [40]. Indeed, a higher prevalence of dog blood parasite infection was observed in dogs from upland areas (villages 4, 5, 6 and 7) compared to lowland areas [39].

Among the 40 positive micromammal samples, we successfully sequenced 11 *Leptospira* amplicons, identifying three species: *L. interrogans* (n = 5; one *Bandicota indica*, three *Berylmys* spp., one *Hylomys suillus*), *L. borgpetersenii* (n = 2 in *Mus* spp.) and *L. weilii* (n = 4; two *Bandicota indica* and two *Berylmys* spp.). While *L. weilii* has been previously identified in rodents in Malaysia [41] and in Papua New Guinea [42], our study represents the first detection of *L. weilii* in rodents in Thailand. A recent study in Nan province (which did not investigate rodents) identified *L. weilii* from cattle, pigs, dogs, and a goat [9]. Outside of Thailand, *L. weilii* has been predominantly detected in cattle and pigs [43,44], but has also been identified in various wildlife species, including bandicoots and grey kangaroos in Australia [44,45] suggesting a broad host range which may vary between countries. In Nan province, *L. interrogans* and *L. weilii* DNA were previously identified in environmental water samples [9], suggesting that transmission between animals and/or species may be enhanced around shared natural water sources and common grazing areas [46].

Our phylogenetic analyses revealed that terrestrial micromammals in Thailand harbor several pathogenic *Leptospira* but they may not contribute equally in all cycles of human infection. *L. interrogans* and *L. borgpetersenii* commonly circulate in humans, dogs and micromammals. *L. weilii* was found in humans, livestock, and dogs but it might be less common in micromammals in Thailand, which could suggest a different transmission cycle for this species. Domestic animals like dogs may act as ‘bridge’ species, epidemiologically connecting humans and unexplored wildlife species [47]. *L. kirschneri* and *L. wolffii* have both been identified in human clinical cases in Thailand, but were not identified in our study in Nan province. In previous animal studies in Thailand, *L. kirschneri* has been detected in one *Mus cervicolor* sampled in Loei [7], but the *rrs* sequence was not published, while *L. wolffii* has been identified in one *Bandicota indica* sampled in Buriram [8]. When blasting the *L. kirschneri* sequences from Thai human clinical cases, the closest sequences from GenBank were from humans (Bulgaria, 99.8% identity; Sri Lanka, 99.6% identity), and cattle and capybara (*Hydrochoerus hydrochaeris*) from Brazil (99.6% identity). This may suggest that, in Thailand, *L. kirschneri* is associated with underexplored animal hosts or ecological niches. This knowledge gap limits our understanding of the complete transmission cycle and zoonotic sources of *L. kirschneri* and *L. wolffii* infections in Thailand, thus underscoring the importance of more integrated molecular surveillance across different animal hosts, including domestic and wildlife species, using common sequencing targets such as *secY* gene.

## Conclusions

The present study demonstrates a complex and dynamic epidemiological picture of *Leptospira* infection in rural Nan Province in Northern Thailand, where dogs and/or multiple terrestrial micromammal species may play a role. In micromammals, we identified *L. interrogans, L. weilii*, and *L. borgpetersenii* but host species varied, which may indicate some level of host specificity of the pathogen. In our study, free-roaming dogs were likely to visit contaminated sites, making them potential bridge hosts for zoonotic transmission. This role could be further amplified by risky behaviors in dogs, such as frequent contact with natural water sources. In such settings, routine dog vaccination could be a relevant public health intervention strategy. However, vaccine protection will depend on local serovar distribution, as commercial canine vaccines may not protect against all circulating pathogenic serovars in an area. Our results underscore the importance of increased molecular surveillance across wildlife and domestic hosts including livestock for comprehensive phylogenetic analyses, which can enhance the understanding of local epidemiology and the identification of animal reservoirs and risk factors. In the future, multidisciplinary approaches—such as improved diagnostic, serological and epidemiological surveys, social and ecological analyses, including behavioral and spatial surveys—can provide a clearer understanding of transmission dynamics and infection hotspots. Only then can we offer tailored public health interventions to break transmission cycles and lower leptospirosis burden in Thai rural communities.

## Acknowledgements

We thank all the dog owners who allowed serum sampling to complete the study, as well as contributed to the questionnaire survey. We especially thank the subdistrict Administration Office (SAO), the village chiefs and volunteers of Saen Thong for their contribution and help.

## Supporting Information

**S1 Table.**
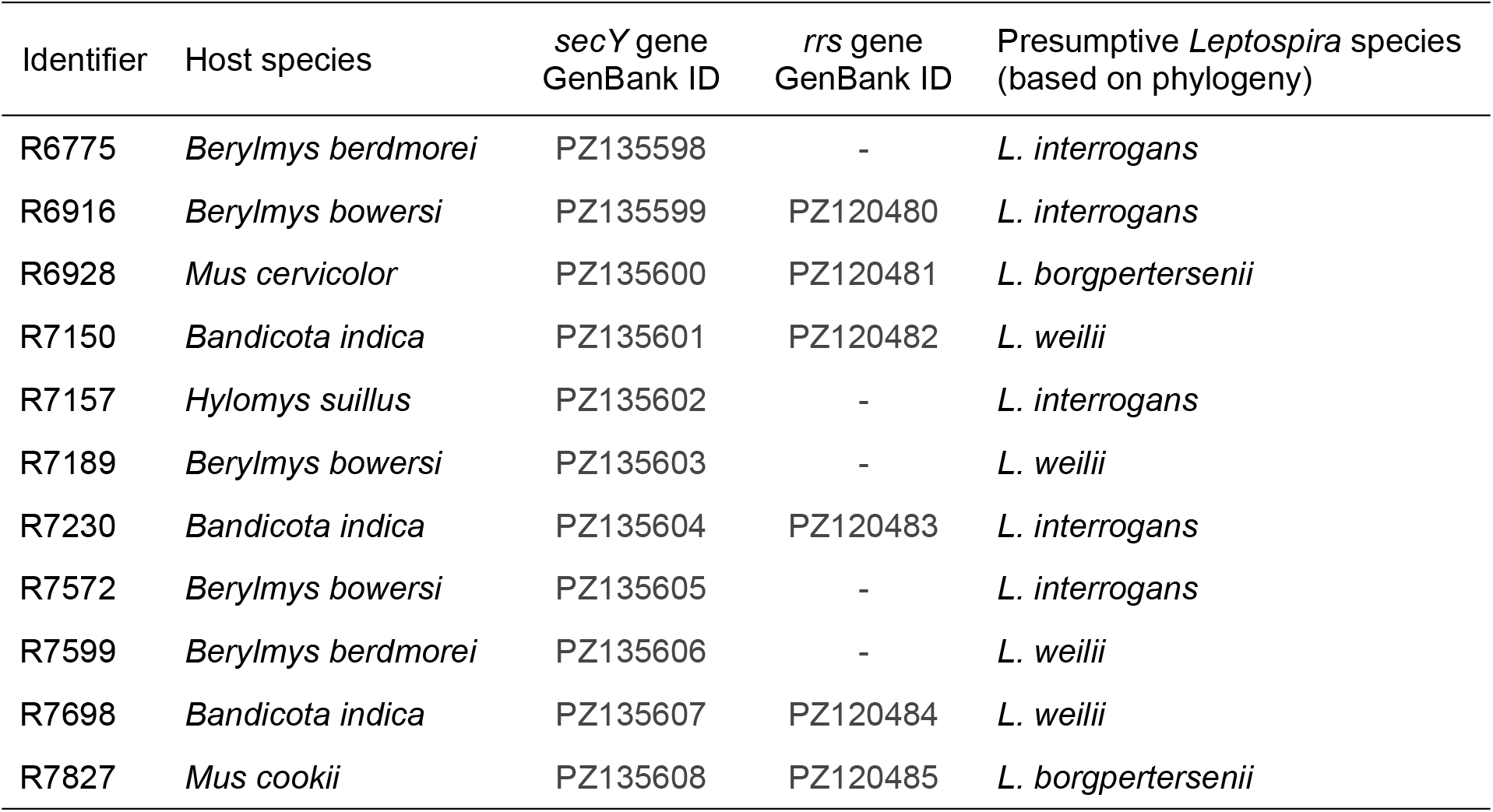
GenBank accession numbers of our *Leptospira* sequences. Sequences for *secY* gene and *rrs* (16S) gene are provided.

**S2 Table.** *Leptospira* infection results per village. N: number of terrestrial micromammals trapped per village. *Lborg: Leptospira borgpetersenii; Lint: Leptospira interrogans; Lwei: Leptospira weilii*; Lspp: unidentified *Leptospira* species. The prevalence of *Leptospira*-positive micromammals per village is provided, as well as the distribution of the 40 qPCR-positive micromammals between villages.

| Village | N | *Lborg* | *Lint* | *Lwei* | Lspp. | Lepto+/40 (%) | Prevalence (%) |
| --- | --- | --- | --- | --- | --- | --- | --- |
| 1 | 8 | 0 | 0 | 0 | 0 | 0/40 (0) | 0/8 (0) |
| 2 | 12 | 0 | 0 | 0 | 1 | 1/40 (2.5) | 1/12 (8.3) |
| 3 | 24 | 0 | 0 | 0 | 1 | 1/40 (2.5) | 1/24 (4.2) |
| 4 | 8 | 0 | 0 | 0 | 0 | 0/40 (0) | 0/8 (0) |
| 5 | 30 | 0 | 0 | 0 | 2 | 2/40 (5) | 2/30 (6.7) |
| 6 | 288 | 1 | 4 | 4 | 22 | 31/40 (77.5) | 31/288 (10.8) |
| 7 | 18 | 1 | 1 | 0 | 1 | 3/40 (7.5) | 3/18 (16.7) |
| 8 | 11 | 0 | 0 | 0 | 2 | 2/40 (5) | 2/11 (18.2) |
| Total | 399 | 2 | 5 | 4 | 29 | 40/40 (100) | 40/399 (10) |

**S3 Table.** Results of the binomial generalized linear mixed model testing the effects of environmental and host factors on *Leptospira* infection in terrestrial micromammals. Body masses were standardized to a species-specific Z-score. Random intercepts accounted for year and village * year interaction. Analyses were limited to species with at least five individuals.

| Predictor | Estimate | Standard error | z-value | p-value |
| --- | --- | --- | --- | --- |
| Intercept | -2.628 | 0.988 | -2.659 | 0.008 |
| Landscape: Lowland | 1.077 | 0.809 | 1.332 | 0.183 |
| Landscape: Settlement | 0.283 | 1.181 | 0.240 | 0.811 |
| Landscape: Upland | 0.520 | 0.720 | 0.723 | 0.470 |
| Season: Wet | -0.348 | 0.623 | -0.559 | 0.576 |
| Sex: Male | -0.336 | 0.409 | -0.821 | 0.412 |
| Standardized body mass | 0.430 | 0.195 | 2.199 | 0.028 |
| Species: *Berylmys berdmorei* | 1.417 | 0.992 | 1.428 | 0.153 |
| Species: *Berylmys bowersi* | 1.008 | 0.867 | 1.162 | 0.245 |
| Species: *Hylomys suillus* | 0.846 | 1.358 | 0.624 | 0.533 |
| Species: *Maxomys surifer* | -0.298 | 1.084 | -0.275 | 0.784 |
| Species: *Mus caroli*<sup>a</sup> | -18.017 | 9125.187 | -0.002 | 0.998 |
| Species: *Mus cervicolor* | 0.047 | 0.704 | 0.066 | 0.947 |
| Species: *Mus cookii* | 0.599 | 0.874 | 0.685 | 0.493 |
| Species: *Niviventer mekongis* | -0.979 | 1.299 | -0.754 | 0.451 |
| Species: *Rattus andamanensis* | -0.021 | 1.349 | -0.016 | 0.987 |
| Species: *Rattus exulans* | 0.068 | 1.132 | 0.060 | 0.952 |
| Species: *Rattus tanezumi* | -1.186 | 1.190 | -0.997 | 0.319 |
| Randoms effects | Variance | Standard deviation |  |  |
| Year | 0.000 | 0.000 |  |  |
| Village * year | 0.118 | 0.343 |  |  |

<sup>a</sup> No *Leptospira*-positive individual was detected in *Mus caroli*, resulting in unstable coefficient estimates due to complete separation. This species-level estimate should therefore be interpreted with caution.

**S1 Figure.**
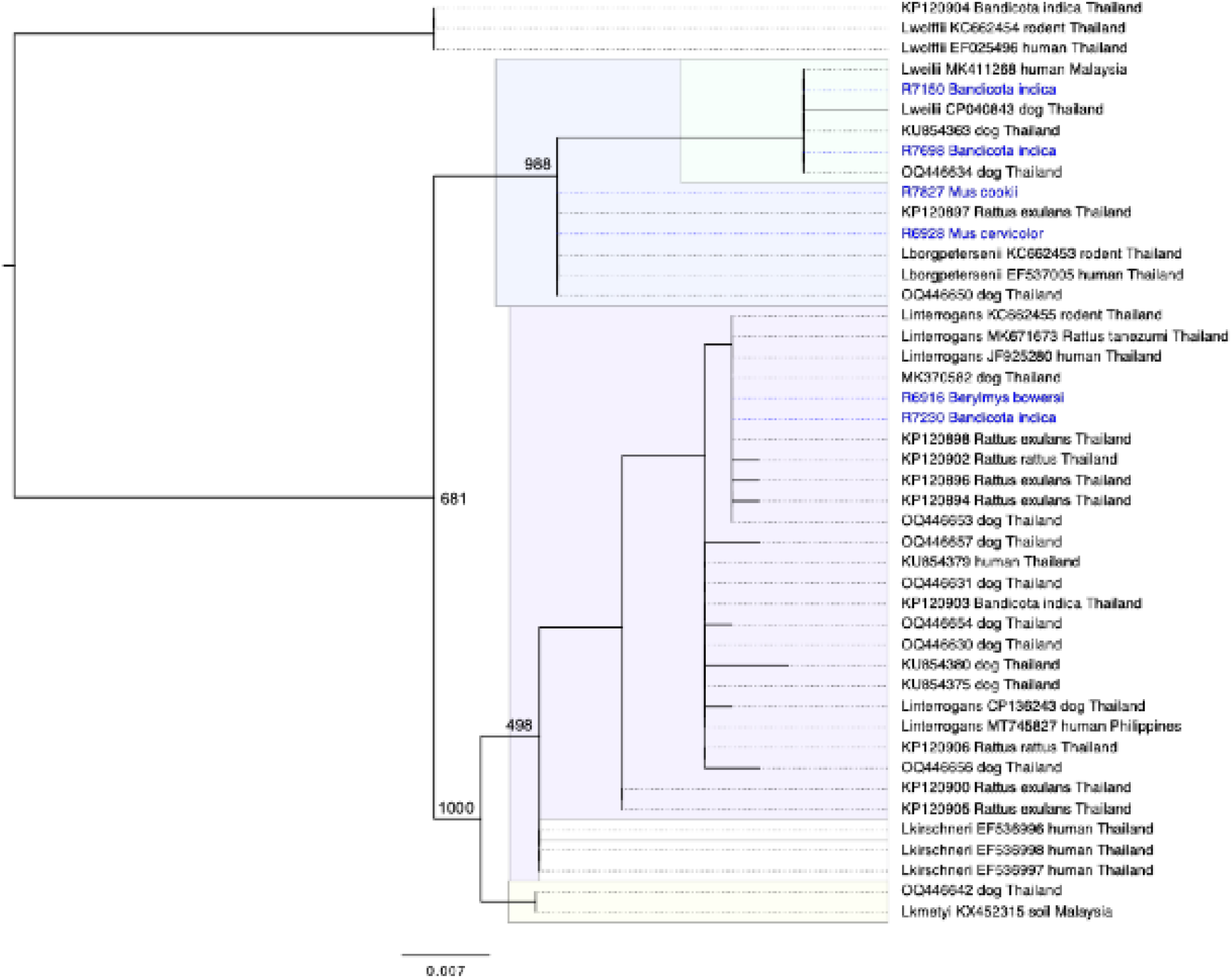
Maximum-likelihood phylogenetic tree (model HKY85; 1,000 replicates) inferred from *rrs* gene (429-bp sequence). Samples from our study (n = 6) are in blue, using identifiers accompanied by the host species. Published sequences (in black) are identified using *Leptospira* species followed by the GenBank accession numbers, the source/host species and the country of origin. Bootstrap values are indicated for the main nodes.

